# Global, Low Amplitude Cortical State Predicts Response Outcomes in a Selective Detection Task

**DOI:** 10.1101/2021.07.23.453568

**Authors:** Krista Marrero, Krithiga Aruljothi, Behzad Zareian, Chengchun Gao, Zhaoran Zhang, Edward Zagha

## Abstract

Spontaneous neuronal activity strongly impacts stimulus encoding and behavioral responses. We sought to determine the effects of neocortical prestimulus activity on stimulus detection. We trained mice in a selective whisker detection task, in which they learned to respond (lick) to target stimuli in one whisker field and ignore distractor stimuli in the contralateral whisker field. During expert task performance, we used widefield Ca^2+^ imaging to assess prestimulus and post-stimulus neuronal activity broadly across frontal and parietal cortices. We found that lower prestimulus activity correlated with enhanced stimulus detection: lower prestimulus activity predicted response versus no response outcomes and faster reaction times. The activity predictive of trial outcome was distributed through dorsal neocortex, rather than being restricted to whisker or licking regions. Using principal component analysis, we demonstrate that response trials are associated with a distinct and less variable prestimulus neuronal subspace. For single units, prestimulus choice probability was weak yet distributed broadly, with lower than chance choice probability correlating with stronger sensory and motor encoding. These findings support a low amplitude, low variability, optimal prestimulus cortical state for stimulus detection that presents globally and predicts response outcomes for both target and distractor stimuli.

## Introduction

The brain is never silent. Throughout sleep and wakefulness, spontaneous neuronal activity reflects dynamic, self-organized states that affect the generation and propagation of neuronal signals (Arieli et al., 1995; Arieli et al., 1996; Ferezou et al., 2007; McCormick et al., 2015; McGinley, David, et al., 2015; McGinley, Vinck, et al., 2015; Niell & Stryker, 2010; Poulet et al., 2012; Zagha & McCormick, 2014). Changes in spontaneous activity impact the amplitude of neuronal sensory responses (Crochet & Petersen, 2006; Haider & McCormick, 2009; Poulet & Petersen, 2008; Sachdev et al., 2004; Shimaoka et al., 2018) and behavioral outcomes (Boly et al., 2007; Fiebelkorn & Kastner, 2021; Kim & Sejnowski, 2021; Mazaheri et al., 2011; McGinley, David, et al., 2015; van Kempen et al., 2020). In awake subjects, these changes correlate with changes in task engagement, movement, and internal (cognitive or egocentric) versus external (perceptive or allocentric) processing modes (Andreou & Borgwardt, 2020; Boly et al., 2007; de Lange et al., 2013; Murphy et al., 2018; Musall et al., 2020; Salkoff et al., 2020; Stringer et al., 2019). However, most studies of sensory processing and sensory detection normalize post-stimulus by prestimulus activity, thereby obscuring the impacts of spontaneous activity. And yet, understanding how spontaneous activity impacts neuronal signaling and task performance will reveal important principles of context-dependent sensory and motor processing.

This study focuses on prestimulus activity during a sensory detection task, for which many open questions remain. First, is the ability to detect a stimulus improved by high or low prestimulus activity (Figure 1A)? A common model of decision-making is integration to bound, which proposes that a decision is made once neuronal activity reaches a specific threshold (Gold & Shadlen, 2007; Hanes & Schall, 1996; Roitman & Shadlen, 2002). Within this model, higher prestimulus activity may bring a network closer to decision threshold and/or increase the gain of a network and therefore promote stimulus detection (Haider & McCormick, 2009). Consistent with this framework, studies in primary visual cortex demonstrate that higher prestimulus activity leads to larger amplitude stimulus responses (Haider et al., 2007). However, higher prestimulus activity may reduce cortical sensory responses (Hasenstaub et al., 2007), due to increased cortical inhibition and reduced intrinsic and synaptic excitability. Studies in the primary somatosensory and primary auditory cortices support this alternative noise suppression framework, demonstrating that lower prestimulus activity, or activity in a low-arousal synchronized state, leads to larger amplitude stimulus responses (McGinley, David, et al., 2015; Petersen et al., 2003; Sachdev et al., 2004).

**Figure 1:**
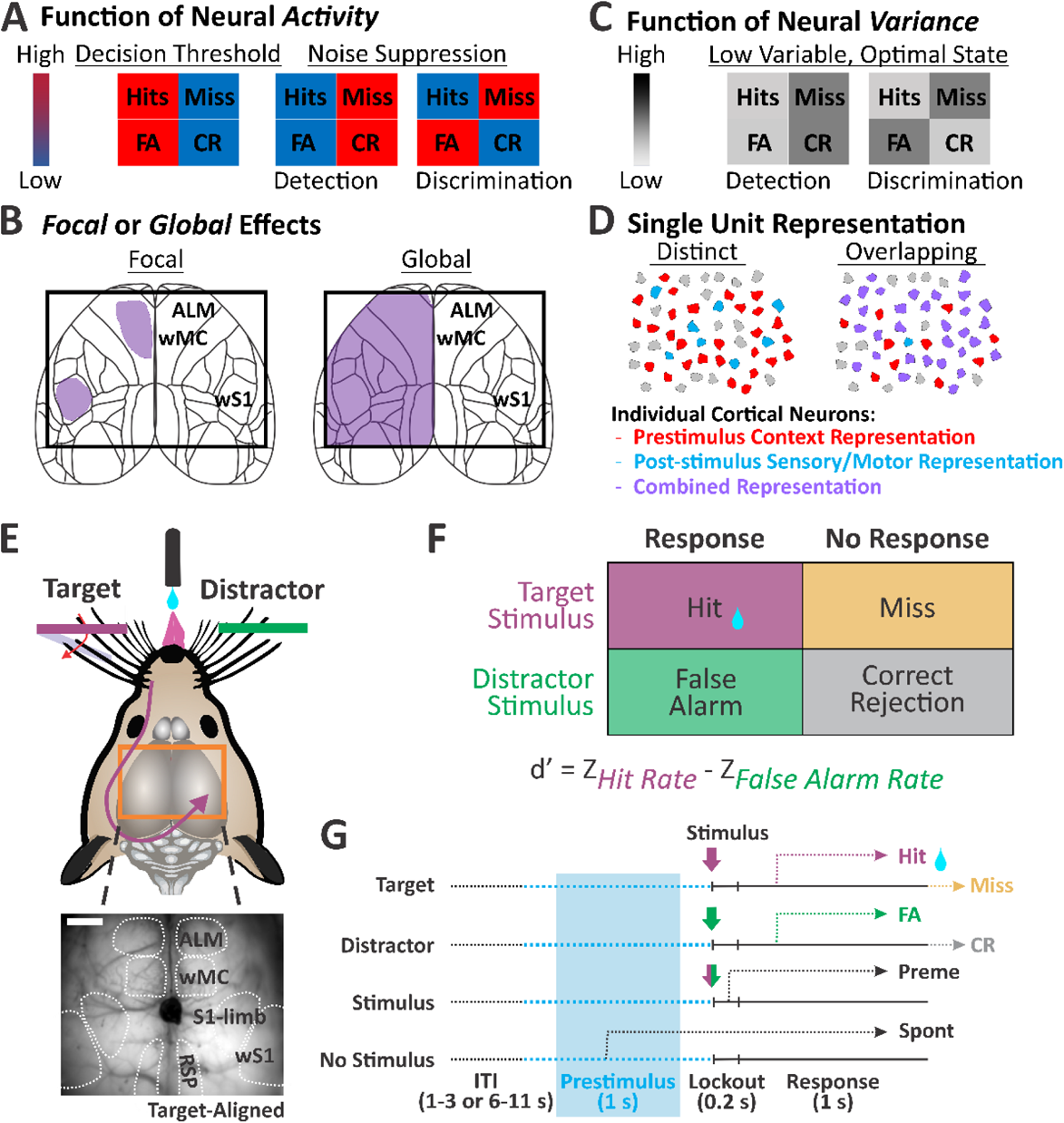
Predictions and experimental design for testing impacts of prestimulus activity on sensory detection and discrimination. (A-D) Potential mechanisms of task-relevant prestimulus activity. (E-G) Experimental design. (E). Head-fixed mice are trained to discriminate between target whisker deflections (purple) and distractor whisker deflections (green), within opposite whisker fields. Mice report detection by licking a central lickport. The orange rectangle reflects the widefield Ca^2+^ imaging window. The inset below is a sample imaging frame, demarcating neocortical regions of interest in bilateral frontal and parietal cortices. (F) Classification of trial types and outcomes. Task performance is quantified by discrimination d’ as the separation between hit and false alarm rates. z, inverse cumulative function of the normal distribution. G. Trial structure, including a variable inter-trial interval, 1 s prestimulus window, 0.2 s stimulus and lockout (delay) window, and 1 s response window. The prestimulus window of interest in this study is the last 1 s of the inter-trial interval (blue shade), immediately before stimulus onset. Spont, spontaneous responses during the prestimulus window; Preme, premature responses during the lockout window. Scale bar in (E) is 1 mm.

In somatosensory (whisker) detection tasks, impacts of prestimulus activity on stimulus encoding and detection have been studied at the level of membrane potential through whole cell patch clamp recordings. While prestimulus membrane potential activity of primary somatosensory cortical neurons did predict sensory response amplitudes (Sachidhanandam et al., 2013), it did not predict trial outcome (e.g., hit versus miss) (Sachidhanandam et al., 2013; Yang et al., 2016). However, these whole cell recording studies are limited by relatively small samples sizes (10s of neurons) which may limit the ability to resolve small yet widespread contributions of prestimulus activity to task performance.

A second open question is whether the prestimulus activity that impacts stimulus encoding and detection is focal and restricted to specific cortical regions or global and observed throughout neocortex (Figure 1B). Global activity may reflect changes in arousal and movement (Musall et al., 2020; Salkoff et al., 2020; Stringer et al., 2019) whereas focal changes may reflect shifts in, for example, attentional focus or response preparation (Fries et al., 2001; Ghose & Maunsell, 2002; Luck et al., 1997; Moore & Armstrong, 2003). It is currently unknown whether prestimulus activity in sensory compared to motor cortices have larger impacts on task performance, and whether the directionality of that impact is the same across neocortical regions (Shimaoka et al., 2018). In addition to considering different cortices individually, is there an ‘optimal state’ of prestimulus activity that includes the contributions of multiple cortices (Figure 1C)? A third open question is whether prestimulus activity has the same or different impacts on target (attended) versus distractor (unattended) stimulus encoding and detection (Figure 1A, C). For example, the same prestimulus activity may promote discrimination (response to targets, no response to distractors) or bias responses for detection (respond to or ignore all stimuli). Lastly, do the neurons that express task-relevant changes in prestimulus activity overlap with or are they distinct from the neuronal populations that express strong post-stimulus sensory and/or motor activity (Figure 1D)?

We address these questions in the context of a selective whisker detection task in mice. We trained mice to respond (lick) to deflections on one whisker field (target) and ignore deflections in the contralateral whisker field (distractor) (Aruljothi et al., 2020; Zareian et al., 2021). Using widefield Ca^2+^ imaging, we previously identified the cortical regions that are highly active post-stimulus and pre-response, and therefore may contribute to stimulus detection: the whisker region of primary somatosensory cortex (S1), the whisker region of primary motor cortex (wMC), and the pre-motor licking region anterior lateral motor cortex (ALM) (Aruljothi et al., 2020). We consider these cortical regions to be ‘task-related’ and all other cortical regions to be ‘task-unrelated’. Here, we implement a sliding window normalization to preserve prestimulus fluctuations. We investigate the impacts of prestimulus activity levels on trial outcome, for both target and distractor stimuli. Additionally, we use dimensionality reduction of the imaging data to assess prestimulus variability across cortices. Lastly, we assess prestimulus choice probability of single units in task-related cortices to determine the distribution of these signals across the neuronal population.

## Methods

The experimental datasets in this study were previously published, including the whisker monitoring, widefield GCaMP6 imaging (Aruljothi et al., 2020) and single unit recordings (Zareian et al., 2021). Below, we summarize these experimental methods and describe the new analyses used in this study.

### Animal Subjects

Experiments were approved by the IACUC of University of California, Riverside. Both male and female adult mice were used, either wild type (C57BL/6J, BALB/cByJ) or transgenic (Snap25-2A-GCaMP6s-D, backcrossed to BALB/cByJ). GCaMP6s expressing transgenic mice were used for widefield Ca^2+^ imaging; wild type mice were used for whisker imaging and electrophysiology. Mice were housed in a 12-hour light/dark cycle; experiments were conducted during the light cycle.

### Animal Surgery

For headpost implantation, mice were placed under isoflurane (1-2%), ketamine (100 mg/kg), and xylazine (10 mg/kg) anesthesia. The scalp was cut (10 mm × 10 mm) and resected to expose the skull. A lightweight metal headpost was fixed onto the skull using cyanoacrylate glue. An 8 mm × 8 mm headpost window exposed most of dorsal cortex. The skull was covered with a thin layer of cyanoacrylate gap-filling medium (Insta-Cure, Bob Smith Industries) to seal the exposed skull and enhance skull transparency; the window was sealed with a quick-dry silicone gel (Reynolds Advanced Materials). Mice were administered meloxicam (0.3 mg/kg) and enrofloxacin (5 mg/kg) for three days post-op. Water restriction began after recovery from surgery (minimum of three days). Training on the behavior rig began after one day of water restriction. For electrophysiological recordings, craniotomies and durotomies (< 0.5 mm diameter) were performed under isoflurane anesthesia. Full recovery from anesthesia was allowed (up to 60 minutes) before placement on the behavioral rig.

### Animal Behavior

Training stages, metrics of learning, and criterion for expert performance in the Go/NoGo selective whisker detection task were previously reported (Aruljothi et al., 2020). Briefly, head-fixed and water deprived mice were placed on a behavioral apparatus controlled by Arduino and custom MATLAB (MathWorks) scripts. Two paddles were placed in whisker fields on the opposite sides of the face, designated as target or distractor. Target and distractor designations were assigned at the beginning of training and remained constant. Following variable intertrial intervals, mice could receive a target trial (rapid deflection of the target paddle), distractor trial (rapid deflection of the distractor paddle) or catch trial (no whisker stimulus). Mice responded by licking at a central lick port. Hits (responses to target stimuli) were rewarded with ∼5 μL of water, correction rejections (not responding to distractor stimuli) and correct withholdings (not responding during the catch trial) were rewarded with a shortened intertrial-interval (ITI) and a subsequent target trial. Licking during the ITI was punished by resetting the ITI, effectively a time-out. Mice were considered expert once they achieved a discriminability *d’* > 1 (separation of hit and false alarm response rates) for three consecutive days:

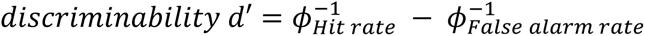

All recordings were conducted in expert mice while performing the task.

### Widefield Imaging

Widefield imaging during expert task performance was conducted as previously reported. The dataset consists of 38 behavioral/imaging sessions, recorded from 5 mice. The through-skull imaging window included bilateral dorsal parietal and frontal cortices. Illumination from a 470 nm LED source (Thorlabs) was band-pass filtered for excitation (Chroma ET480/40x) and directed onto the skull via a dichroic mirror (Chroma T510lpxrxt). Emitted fluorescence was band-pass filtered (Chroma ET535/50m) and collected using an RT sCMOS camera (Diagnostic Imaging, SPOT Imaging software). Images were acquired at 10 Hz with a final resolution of 142 × 170 pixels (41 μm per pixel). Image sequences were imported to MATLAB for subsequent analyses.

### Electrophysiology

Single unit recordings during expert task performance were conducted as previously reported (Zareian et al., 2021). The dataset consists of 32 behavioral/recording sessions, recorded from 22 mice, yielding a total of 936 single units from three cortical regions (target-aligned whisker region of primary somatosensory cortex [S1], whisker region of motor cortex [wMC], and anterior lateral motor cortex [ALM]). Coordinates (mm, from bregma): S1 3.2-3.7 lateral, 1-1.5 posterior; wMC 0.5-1.5 lateral, 1-2 anterior; ALM 1-2 lateral, 2-2.5 anterior. Recordings were targeted to layer 5 of S1, wMC, and ALM, approximately 500 to 1100 μm below the pial surface. Electrophysiological recordings were conducted using a silicon multielectrode probe (NeuroNexus A1x16-Poly2-5mm-50s-177), positioned using a Narishige micro-manipulator. Neuralynx amplifier (DL 4SX 32ch System) and software were used for data acquisition and spike sorting.

### Whisker imaging

Whisker imaging during expert task performance was conducted as previously reported (Aruljothi et al., 2020). The dataset consists of 9 behavioral/recording sessions, recorded from 4 mice. Images were acquired with a CMOS Camera (Thorlabs DCC3240M camera with Edmund Optics lens 33-301) at either 20 or 60 Hz. No systematic difference between 20 and 60 Hz was observed (data not presented). The imaging field of view included both paddles and the mouse’s head (including whiskers and snout).

### Data Analysis

Data analyses were performed in MATLAB using custom scripts.

#### Engagement period

To ensure that analyses were conducted during task engagement, ‘engaged periods’ were defined as continuous behavioral performance of at least 10 minutes without 60 seconds of no responding. For sessions with more than one engaged period, the longest engaged period was used for further analyses. Furthermore, sessions were included in subsequent analyses only if performance was at expert level: discriminability d’>1. For sessions with multiple stimulus amplitudes, trials were combined for further analyses only when the differences in response rates were 15% or less.

#### Sliding Window Normalization and Trial-Based Neuronal Activity

The trial-based imaging time window consisted of the prestimulus epoch (1 s), the stimulus and lockout epoch (0.2 s), and the allowable response epoch (1 s), a total of 2.2 s. A raw movie F was created by concatenating fluorescence activity from consecutive trials, where *F_n_(i,j,f)* is the fluorescence of each pixel (row *i* and column *j*) in frame *f* for each trial *n*. To generate normalized fluorescence values, we first determined the sliding window local mean for each pixel, computed every 2 s using a +/- 200 s window size [*F_SW_(i,j,n)*]. Then, we calculated the normalized fluorescence (Salkoff et al., 2020) (see also Supplemental Figure 1) for each pixel at each frame as:

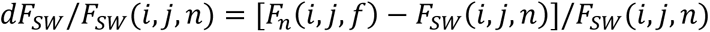

Trialwise average movies were then compiled by first indexing outcome type (*hit*, *miss*, *false alarm*, *correct rejection*) and then by averaging pixelwise activity across corresponding frames of corresponding trials. Frames were aligned to the stimulus onset frame (stimulus-aligned) where stimulus occurred or aligned to the first frame containing the response (response-aligned) where response occurred. Trials with responses during the lockout period were considered *premature* and excluded from the analysis. Trials with responses before the stimulus but within the prestimulus imaging period were considered *spontaneous*, dF/F reported but not further analyzed. Grand average movies were aligned to bregma, flipped at bregma according to target-distractor assignment, and then averaged across all sessions.

#### Difference in Prestimulus Fluorescence

Fluorescence differences for target and distractor assignment were calculated per trial type per session. Prestimulus frames 6 to 10 (capturing the last 500 ms of the prestimulus window, before stimulus onset) were trialwise and pixelwise averaged per session. Session data were excluded from this analysis if there were fewer than 5 incorrect trials in the session (excluding 9 sessions for target Miss, 6 sessions for distractor FA). For target fluorescence difference frames (n = 29 sessions), Hits fluorescence mean frame was subtracted from Miss fluorescence mean frame. For distractor fluorescence difference frames (n = 32 sessions), FA fluorescence mean frame was subtracted from CR fluorescence mean frame. Response prestimulus frames were subtracted from no response prestimulus frames because no response fluorescence activity was generally higher than response fluorescence activity. Prestimulus difference frames were aligned, assigned, and averaged across all sessions (as above). To normalize for differences in changes in fluorescence across regions, we performed a z-score normalization of the dF/F values for each pixel as the pixelwise mean divided by the pixelwise standard deviation (μ_i,j_/σ_i,j_). For quantification of target versus distractor prestimulus difference, normalized difference (index), and significance, frames were averaged across pixels for scalar values.

#### Regression analyses between Prestimulus Activity and Reaction Time for Response Trials

The correlation between activity during prestimulus period (dF/F) and reaction times (RT) for response trials (Hits and FAs) were computed as a linear regression from which we obtained the slope of the linear fit with 95% confidence interval and coefficient of determination, R^2^, as the goodness of fit (Zareian et al., 2020) (Curve Fitting Toolbox in Matlab). For this analysis, we assigned prestimulus dF/F as the independent variable and reaction time as the dependent variable.

#### Stimulus Encoding in Post-Stimulus Fluorescence

Stimulus encoding was quantified as the neurometric *d’* (Britten et al., 1992) of prestimulus fluorescence (stimulus absent) and post-stimulus fluorescence (stimulus present) during the lockout epoch, as previously applied to imaging data (Aruljothi et al., 2020). We excluded session data from this analysis if there were fewer than 4 incorrect trials in the session (excluding 5 sessions for target Miss, 2 sessions for distractor FA). Neurometric *d’* was calculated separately according to target and distractor assignment and then according to trial type outcome. This resulted in 6 different datasets for stimulus encoding: all target, all distractor, hit trials, miss trials, false alarm trials, and correct rejection trials. Prestimulus and post-stimulus fluorescence histograms were plotted into receiver operating characteristic (ROC) curves and the area under the curve (AUC) was converted to *d’* as the neurometric:

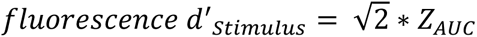

Region specific pixel values were identified as the maximum value within the defined regions of interest (ROI), performed for target-aligned and distractor-aligned regions of S1, wMC, and ALM. The difference in stimulus encoding in S1 between the response and the no response trials for both target and distractor stimuli was calculated as the percentage:

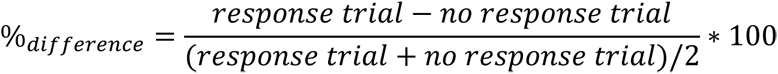

#### Whisker Motion Energy During Behavior

The imaging window was cropped by region of interest: target or distractor paddle stimulus or whisker fields. The function *vision.VideoFileReader* was used for optimal reading of video frames into MATLAB. Whisker movement per frame (*Δframe*) was calculated as the pixelwise frame by frame mean gray value (*MGV*) difference (*ΔMGV_pixel_*). Whisker motion energy (*WME*) was defined as the sum of the squares across pixels:

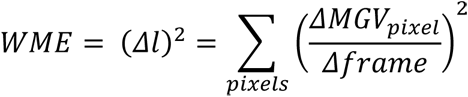

WME traces of the cropped videos of the paddles were used to detect stimulus events (target/distractor). This was performed by using a constant threshold and aligning detected events from the video to their temporally closest events recorded using Arduino. The traces from the cropped videos of whisker fields were transformed (z-scored) to have a mean of zero and standard deviation of 1 for the purpose of comparison across sessions. Subsequently, WME data were temporally aligned by trial type to stimulus onset (target/distractor) determined from the videos.

#### Principal Component Analysis of Fluorescence

Fluorescence was averaged across anatomic masks [target and distractor S1, wMC, ALM, and retrosplenial (RSP) cortex] per frame per trial per session. Mean regions were normalized and placed into a covariance matrix. The covariance matrix was decomposed into an eigenmatrix, eigenvectors were sorted by eigenvalue weight, and eigenvectors were projected into component space. All frames were separated by trial type, plotted in PC space, and differentiated by trial epoch (prestimulus, post-stimulus and pre-response lockout, and allowable response window). Component data for prestimulus frames were further analyzed: confidence area ellipses of 1 standard deviation, σ, was defined by the ellipsoid distribution of prestimulus frames in PC space per session. Centroids were defined as the geometric mean of prestimulus frames in PC space per session.

#### Spike Sort and Cluster of Single Units

Using Neuronalynx recording system, signals were sampled at 32 kHz, band-pass filtered from 0.1Hz to 8000 Hz, and high-pass filtered at 600 Hz to 6000 Hz. Putative spikes crossed thresholds of 20 to 40 μV, isolated from baseline noise. KlustaKwik algorithm in SpikeSort3D software was used for spike sorting and clustering. Clusters were defined by waveform and cluster location in feature space (peaks and valleys); movement artifacts (atypical waveforms or those occurring across all channels) were removed, as previously reported (Zareian et al., 2021). Subsequent analyses were conducted using MATLAB software (MathWorks).

#### Sensory and Motor Encoding of Single Units

Sensory and motor encoding of single units was performed as previously reported (Zareian et al., 2021). Sensory encoding was quantified by the neurometric *d’* using stimulus absent spiking (300 ms prestimulus) and stimulus present spiking (100 ms post-stimulus). Motor encoding was quantified by the neurometric *d’* using response absent spiking (300 ms prestimulus) and response present spiking (100 ms pre-response). Distributions were plotted into ROC curves and the AUC was converted to *d’* as a neurometric:

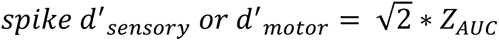

#### Choice Probability of Single Units

For choice probability analyses, we ensured that there was a minimum of 5 trials per trial type (minimum 5 Hits and 5 Miss). Choice probability (%) was quantified as the separation of prestimulus spiking in Hits versus Miss trials. ROC and AUC were calculated from the distributions of Hits and Miss across trials, 500 ms to 0 ms before stimulus onset, averaged over 50 ms nonoverlapping intervals, as previously reported (Zareian et al., 2021)..

#### Statistical Analyses

For imaging statistics, threshold for statistical significance was corrected for multiple comparisons with a Bonferroni correction. Fluorescence difference (Miss – Hits, CR – FA), statistical analyses determined whether dF/F frames were significantly different than zero across sessions (one sample t-test). For whisker analyses statistics, since the number of samples in the whisking data were low, we used one-sample Kolmogorov-Smirnov test (*kstest* in MATLAB) to test for normality assumptions. Since the data mostly violated the normality assumption, Wilcoxon signed rank (*signrank* in MATLAB) and rank sum (*ranksum* in MATLAB) tests were used for comparisons between prestimulus and post-stimulus whisking and between trial types (Hits vs. Miss, FA vs. CR), respectively. For stimulus encoding (neurometric d’), statistical analyses determined whether the trialwise (Hits, Miss, FA, CR) maximum pixel value in S1 was significantly different than zero across sessions (one sample t-test). For differences in stimulus encoding, statistical analyses determined whether the stimulus-aligned S1 maximum pixel value was significantly different between response (Hits, FA) and no response (Miss, CR) outcome types across sessions (two sample t-test). For PCA ellipsoid variance and centroid distribution, statistical analysis determined whether ellipsoid variance or centroid distribution was significantly different between response and no response prestimulus frames, evaluated per component. Box whisker plots show the distribution of prestimulus frames or ellipsoid centroids per trial type with outliers, evaluated per component. For choice probability of single units, statistical analysis determined whether distributions within regions were significantly different from chance (one-sample t-test, chance level 50%) and whether distributions between regions were significantly different from each other (ANOVA and post-hoc Tukey test). For the significance assessment of sensory and motor encoding of single units, one-sample t-test was used to compare d-prime distributions to zero. For the relationship between sensory and motor encoding and choice probability of single units, statistical analysis determined whether regression slopes were significantly different from zero (95% confidence bounds for slopes). Box whisker plots were used to show distributions of sensory encoding, motor encoding, and choice probability of single units evaluated within regions. Average data are presented as mean +/- standard error of the mean, unless otherwise indicated.

## Results

### Global prestimulus activity predicts response outcomes

We considered how prestimulus activity may influence sensory detection (Figure 1A-D). High prestimulus activity may promote detection of target and distractor stimuli; alternatively, low prestimulus activity may promote detection of target and distractor stimuli or discrimination of target from distractor stimuli (Figure 1A). The prestimulus activity that influences behavioral outcomes may present focally in specific task-related regions or globally across neocortex (Figure 1B). A low variability, specific ‘optimal state’ configuration may promote stimulus detection or target/distractor discrimination (Figure 1C). At the level of single units, prestimulus contextual signals and post-stimulus sensory and motor signals may be carried by distinct neuronal ensembles (sparse coding) or overlapping neuronal ensembles (dense coding) (Figure 1D). We tested these possibilities in a selective whisker detection task, in which head-fixed mice learn to respond to rapid deflections in one whisker field (target) and ignore identical deflections in the opposite whisker field (distractor) (Figure 1E). In this task, the possible trial outcomes include hit (response to target), miss (no response to target), false alarm (FA, response to distractor), and correct rejection (CR, no response to distractor) (Figure 1F). Prior to each stimulus was a variable inter-trial interval (ITI), in which mice were required to withhold responding or else reset the ITI. The prestimulus epoch we used for analyses is the last 1 second of the ITI immediately prior to stimulus onset (Figure 1G).

We used widefield Ca^2+^ imaging to measure neuronal activity during expert task performance in frontal and parietal cortices, bilaterally (Figure 1F). Our imaging dataset consists of 38 imaging sessions from 5 mice, using a single task-engaged period per session (see Methods). Due to the highly lateralized cortical whisker representation, we could clearly define target-aligned and distractor-aligned cortical regions, contralateral to the side of the whisker stimulus. To preserve activity fluctuations prestimulus, we normalized raw fluorescence activity using a sliding window method (±200 s sliding window, see Methods and Supplemental Figure 1).

In Figure 2 we present grand average fluorescence activity for each trial outcome, aligned to the onsets of both the stimulus and response. In the first column of Figure 2 we show the last prestimulus frame, which is representative of the full prestimulus epoch. We note stark differences in prestimulus activity for different trial outcomes, particularly when comparing hit (Figure 2A) and miss (Figure 2D) trials. We observed lower prestimulus activity for hit versus miss and for FA versus CR trials, indicating that lower prestimulus activity precedes ‘response’ compared to ‘no response’ outcomes. Interestingly, low prestimulus activity appears to be specifically related to stimulus detection rather than response preparation. This is evidenced by higher activity preceding spontaneous responses (Spont, a response during the ITI, Figure 2C) compared to stimulus-related responses (hits and FA, Figures 2A and 2B). The magnitude of the prestimulus differences is large, on the same scale as the post-stimulus activity. Additionally, prestimulus activity suppression preceding response trials appears to be widely distributed throughout dorsal neocortex, rather than being focused on the task-related regions of S1, wMC and ALM.

**Figure 2:**
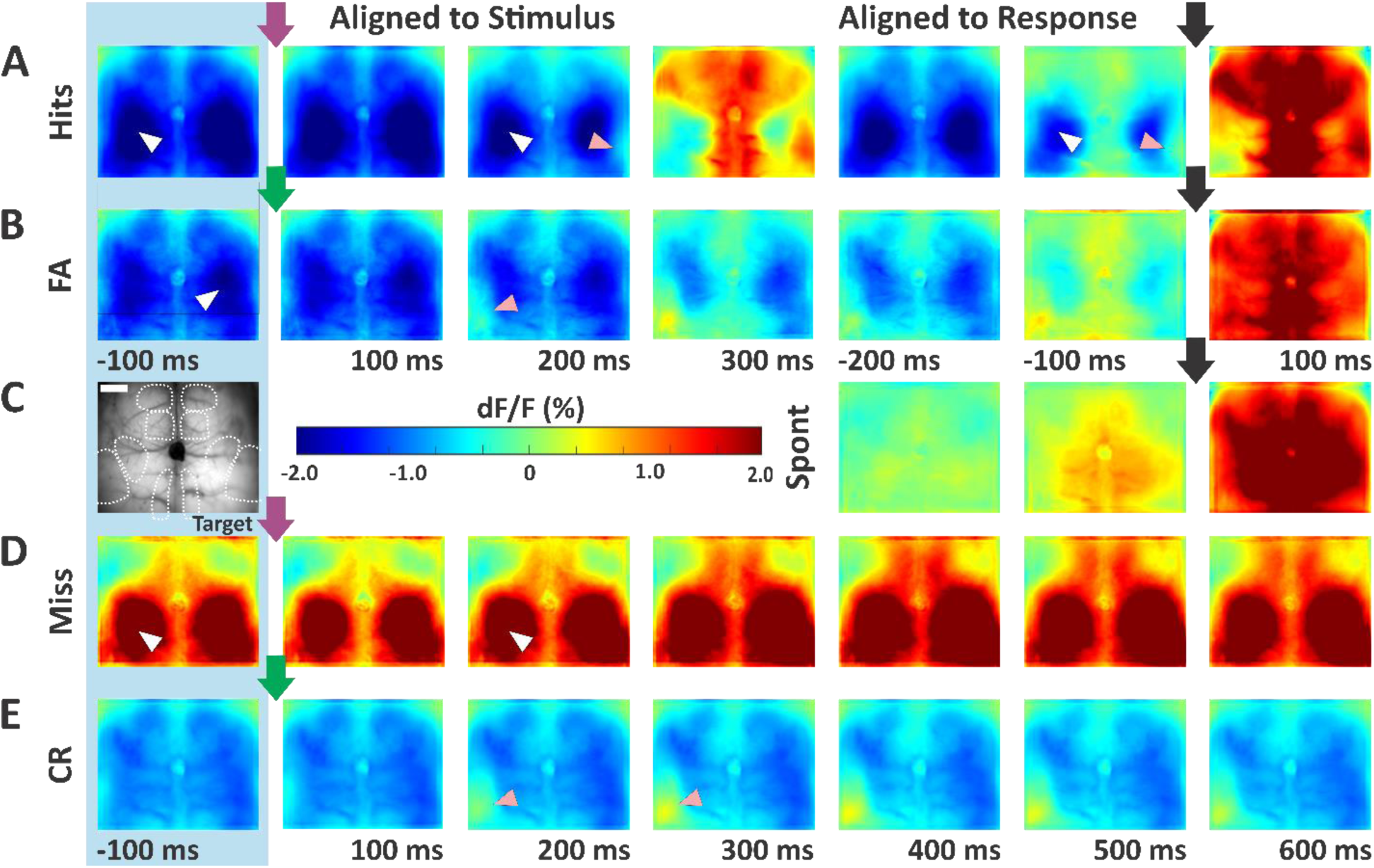
Sliding window normalized grand average fluorescence activity (dF/F). Data are averages across all mice and all sessions (n=38 sessions). Activity in specific imaging frames is aligned to the stimulus onset (left, purple and green arrows for target and distractor stimuli, respectively) or response onset (right, black arrows, in rows A, B, and C). Warmer colors indicate higher activity. The pink arrowheads specify stimulus-aligned whisker regions of S1, whereas the white arrowheads specify limb regions of S1 (see atlas in leftmost panel in row C). The last prestimulus frame is shown in the first column (blue shade). Shown are hit trials (A), false alarm trials (B), spontaneous trials (C), miss trials (D), and correct rejection trials (E). Note the low (negative due to normalization) dF/F prestimulus activity in response trials (hit and false alarm), compared to the high dF/F prestimulus activity in miss trials. Scale bar in (C) is 1 mm.

We quantified the differences in prestimulus activity between response and no response trials for target and distractor stimuli (Figure 3A-F). Shown in this figure are data from the last 500 ms of the prestimulus (similar results were obtained using 100 ms or 1 s prestimulus epochs, data not shown). We subtracted the average prestimulus fluorescence activity of hit from miss trails (Figure 3A). The positive values indicate higher activity preceding miss compared to hit trials (n=29 sessions, averaged across the entire field of view: dF/F μ_[Miss-Hits]_=2.1%±0.3%; one-sample t-test, t(28)=8.1, p=7.9e-09). The largest differences were not in the task-related whisker or licking regions but appear to be focused on the limb regions of somatosensory cortex. While dF/F is already a normalized metric, we sought to further control for possible regional differences in imaging sensitivity. Therefore, we conducted the same subtraction analysis, but on dF/F values that were additionally normalized by z-score, using the mean and standard deviation of the entire session. With this analysis (Figure 3B), the activity differences are more uniformly distributed across frontal and parietal cortices, with an average miss-hit difference of 1.2 standard deviations.

**Figure 3:**
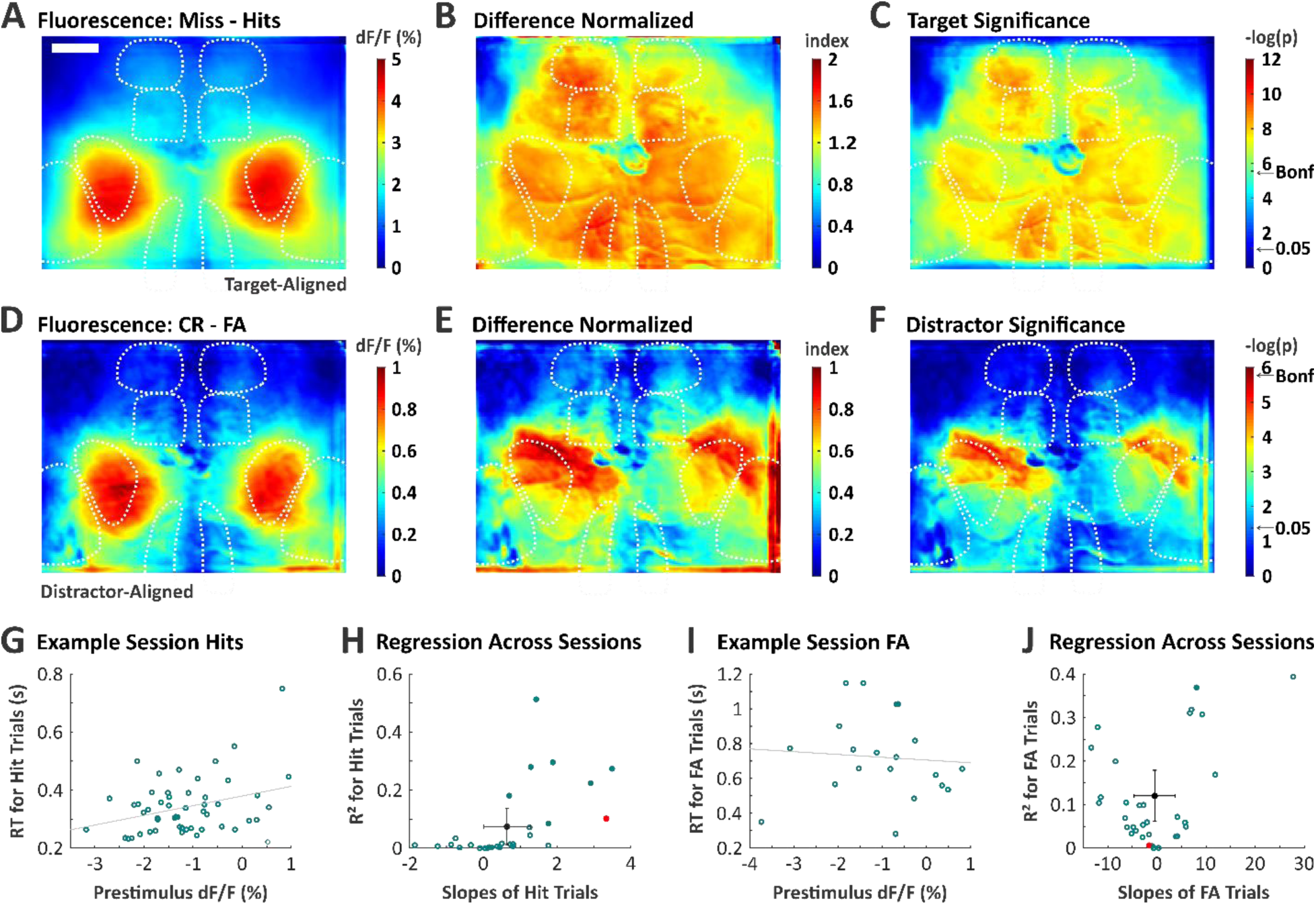
Prestimulus neuronal activity differences between response and no response trials and correlations with reaction time. (A) Grand average of prestimulus dF/F for miss minus hit trials. All pixel values within neocortex are greater than 0, indicating higher global activity preceding miss trials. (B) Similar to [A], except that the individual session dF/F signals were further normalized by z-score to control for differences in fluorescence fluctuations. (C) Significance map for the data in [A]. Significance threshold with Bonferroni correction for multiple comparisons is indicated by the arrow (Bonf). For target trials, higher activity preceding no response trials is statistically significant throughout dorsal cortex. (D-F) Same structure as [A-C], except for CR minus FA trials. Note the more restricted range of scale bars in each panel, compared to target data. For distractor trials, higher activity preceding no response trials is marginally significant, most prominent in the S1 limb regions. Scale bar in (A) is 1 mm. (G) An example session showing a positive correlation between prestimulus activity (dF/F) and reaction time for individual Hit trials (slope=3.34, R^2^=0.10, dotted line is the linear regression). (H) Regression analyses across all sessions for Hit trials. The red data point is the example session in [G], the black data reflect the mean ± standard deviation across sessions (n=30 sessions). (I) FA trials in an example session, with a non-significant negative correlation between prestimulus activity and reaction time (slope=-1.6, R^2^=0.006). (J) Same as H but for FA trials (n=32 sessions).

To determine the spatial regions of significance, on each pixel we performed a paired, two-sample t-test on average prestimulus fluorescence activity in hit versus miss sessions (p-value of each pixel shown in Figure 3C). All neocortical regions within our field of view demonstrated statistical significance, even with a Bonferroni corrected alpha level to control for multiple comparisons (28,960 pixels). Thus, lower prestimulus activity on upcoming target trials is predictive of hit versus miss outcomes. This is observed for all cortical regions within our field of view, including task-related and task-unrelated regions.

There were some notable similarities and differences for distractor trials (Figure 3D-F). Similar to target trials, higher activity was observed preceding no response (CR) versus response (FA) trials (n=32 sessions, averaged across the entire field of view: dF/F μ_[CR-FA]_= 0.36± 0.11% one-sample t-test, t(31)=3.38, p=0.002). However, the fluorescence differences were approximately 5-fold higher for target trials compared with distractor trials (dF/F μ_[Miss-Hits]_=2.1% versus μ_[CR-FA])_=0.36%). A second difference is that for distractor trials, the focus on the somatosensory limb regions was observed in dF/F, z-score, and p-value maps (Figure 3D-F, respectively). The regions with the lowest p-value were slightly above the Bonferroni corrected alpha level. Thus, while lower activity preceding distractor trials was also predictive of a response, the effect size was smaller and less widespread.

In addition to predicting response outcome, we also sought to determine whether prestimulus activity levels predict reaction time on response trials (Figure 3G-J). For these analyses, we determined the slope and coefficient of determination (R^2^) of linear fits for prestimulus dF/F versus reaction time for Hit and FA trials (separately) for each session. As shown in the example session in Figure 3G, a positive slope indicates a correlation between higher prestimulus activity and longer post-stimulus reaction times. Across all sessions, we found a significant positive correlation (positive slope) on Hit trials between prestimulus activity and reaction time (n=30 sessions, slope=0.64±0.23, one-sample t-test: t(29)=2.73, p= 0.011; R^2^=0.074±0.023) (Figure 3H). Thus, for target stimuli, lower prestimulus activity predicts both Hit versus Miss outcomes and faster reaction times.

We performed the same correlation analyses for FA trials (Figure 3I,J). In contrast to Hit trials, FA trials across sessions did not show a consistent correlations between prestimulus activity and reaction time (n=32 sessions, slope=-0.45±1.48, one-sample t-test: t(31)=-0.3, p=0.76; R^2^=0.12±0.021) (Figure 3J).

### Contributions of stimulus encoding and movement on trial outcomes

Next, we assessed whether the differences in trial outcome were reflected in differences in stimulus responses in the neocortex. We quantified the stimulus encoding during the lockout period (200 ms post-stimulus and pre-response) for each trial type (Figure 4). For each pixel, we measured stimulus encoding as the neurometric sensitivity index d’ (Figure 4A-F) and determined whether these values were significantly different from zero (Figure 4G-L). We observed significant stimulus encoding in the stimulus-aligned primary somatosensory cortex (S1) for each trial type (one-sample t-test, n= 38, hits: 38, miss: 33, FA: 36, CR: 38, hits: d’ μ_S1_=0.98±0.06, t(37)=15.58, p=7.79e-18; miss: d’ μ_S1_=0.69±0.08, t(32)=9.08, p=2.26e-10; FA: d’ μ_S1_=1.05±0.09, t(35)=12.08, p=4.87e-14; CR: d’ μ_S1_=0.58±0.049, t(37)= 11.89, p=3.32e-14). Thus, significant stimulus responses occur in S1 for both response and no response trials. However, we did observe a 40-60% reduction in S1 stimulus encoding in no response compared to response trials for target and distractor stimuli (hits vs. miss: d’ μ_% difference_=39.84±7.44%, paired-sample t-test, t(32)= 4.51, p=8.26e-05; FA vs. CR: d’ μ_% difference_=61.62±7.26%, paired-sample t-test, t(35)= 6.72, p=8.75e-08, see Methods). In summary, response trials are associated with reduced prestimulus activity and enhanced post-stimulus sensory responses.

**Figure 4:**
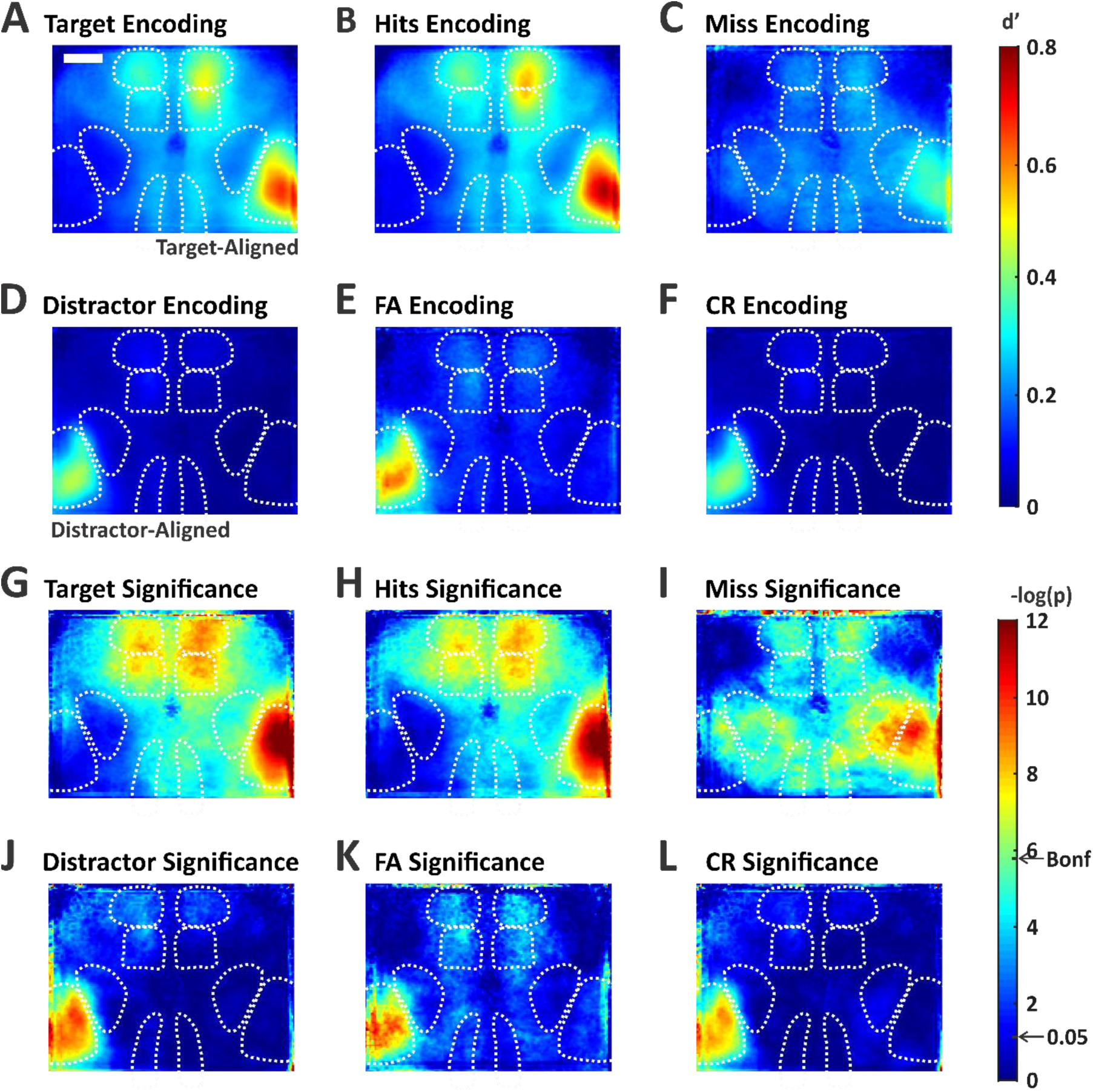
Quantification of stimulus encoding for each trial type. (A-F) Neurometric d’ values were calculated for each pixel during the last frame of the lockout: after stimulus presentation and before the allowed response window. Data are grand average d’ maps from all sessions, showing all target trials (A), hit trials (B), miss trials (C), all distractor trials (D), FA trials (E), and CR trials (F). Note the larger stimulus encoding in response trials (B and E compared to C and F). Significance maps of the data in [A-F], respectively. Significance threshold with Bonferroni correction for multiple comparisons is indicated by the arrow (Bonf). For all trial types there is significant stimulus encoding in the stimulus-aligned S1 whisker region. Scale bar in (A) is 1 mm.

Recent studies have demonstrated widespread neuronal activity increases due to movement (Musall et al., 2020; Salkoff et al., 2020; Stringer et al., 2019). Therefore, in a separate set of recordings, we determined the magnitude of prestimulus and post-stimulus whisker movements on different trial outcomes. Whisker movement was quantified as whisker motion energy (WME, normalized by z-score, see Methods). In Figure 5A-C we present these analyses for one example session for target stimuli. On hit trials, WME increased dramatically post-stimulus (Figure 5A, purple trace). We interpret this as whisking being part of the ‘uninstructed’ behavioral response sequence (Musall et al., 2020). Importantly, we also observed differences in WME prestimulus, with higher WME on miss compared to hit trials (mean +/- STD WME μ_Hits_=-0.45 ± 0.32, WME μ_Miss_=0.19 ± 0.71, rank sum=1516, p=0.001, two-sided Wilcoxon rank sum test; Figure 5A and 5B). In Figure 5C, we show prestimulus WME for each target trial, with the color of the bar indicating trial outcome. High prestimulus WME was more likely to result in a miss trial, even though many miss trials were not preceded by high prestimulus WME. Similar results were observed across all sessions (n=9 session, Figure 5D, Wilcoxon sign rank test, mean +/- STD prestimulus WME μ_Hits_=-0.12 ± 0.17 vs. prestimulus WME μ_Miss_=0.12 ± 0.15, signed rank=1, p=0.008). Thus, high prestimulus movement was associated with some, but not all, of the miss trials.

**Figure 5:**
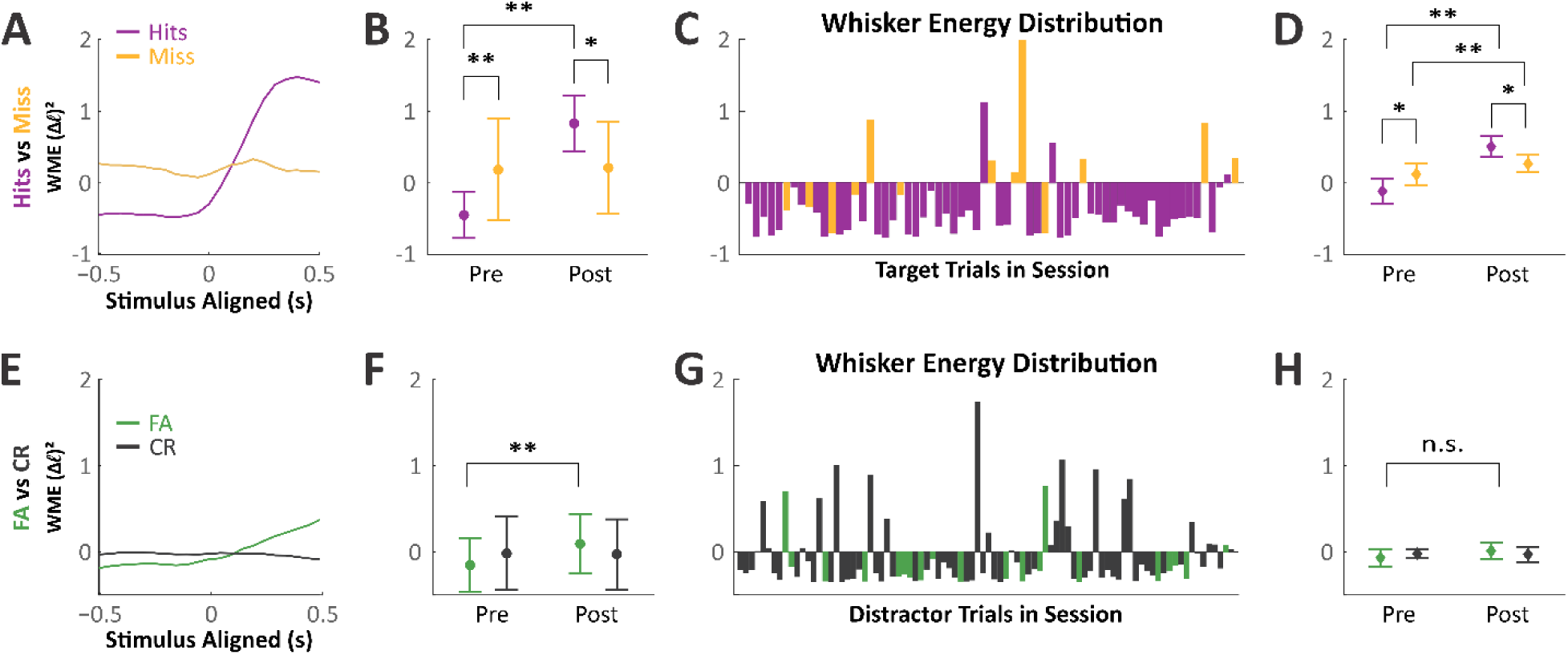
Prestimulus and post-stimulus whisker movements in each trial type. (A) Peristimulus whisker motion energy (WME) on target trials in an example session, hits (purple) and misses (orange). On hits trials there was a dramatic increase in WME post-stimulus and during the response window. Prestimulus, however, WME on hits trials was reduced compared to miss trials. (B) Quantification of data in [A], comparing prestimulus (pre) and post-stimulus WME for hit and miss trials. (C) Prestimulus WME values for each trial in the example session. (D) Summary data for all sessions (n=9). Note the reduced WME preceding hit compared to miss trials. (E-H) Same as above, but for distractor trials. While this example session shows moderately reduced WME preceding false alarm trials (E-G), this trend was not statistically significant across the full dataset (H). Data are presented as mean +/- STD, *p<0.05, **p<0.005.

Differences in prestimulus WME were not as pronounced on distractor trials (Figure 5E-H). We did notice a trend towards increased WME on CR trials. However, this effect was not statistically significant across sessions (n=9 session, Figure 5H, Wilcoxon sign rank test: prestimulus WME μ_FA_=-0.14 ± 0.2 vs. prestimulus WME μ_CR_=-0.04 ± 0.10, signed rank=8, p=0.098). Notably, the effects of prestimulus movement on target and distractor trial outcomes parallel the effects of prestimulus neuronal activity: low prestimulus neuronal activity and low prestimulus WME predict response outcomes, yet these effects are much more pronounced for target compared to distractor trials.

### Analyses of prestimulus activity variance and subspace in reduced spatial dimensions

Next, we sought to characterize frame-by-frame variability in our imaging data. To accomplish this, we used principal component analysis (PCA) to reduce the spatial dimensionality (Figure 6). First, we extracted regional single-trial fluorescence activity using anatomic masks from the dorsal neocortex centered on regions of interest: target/distractor S1, RSP, wMC, and ALM (Figure 6A). We concatenated data from all frames, trials, sessions, and mice and performed PCA on this combined matrix. This enabled us to convert all sessions into the same lower-dimensional axes. Most of the variability in our imaging data could be explained by the first component (∼91%) and the first two components explained ∼96% of the variance (Figure 6B-D). Therefore, further analyses focused on these first two spatial components.

**Figure 6:**
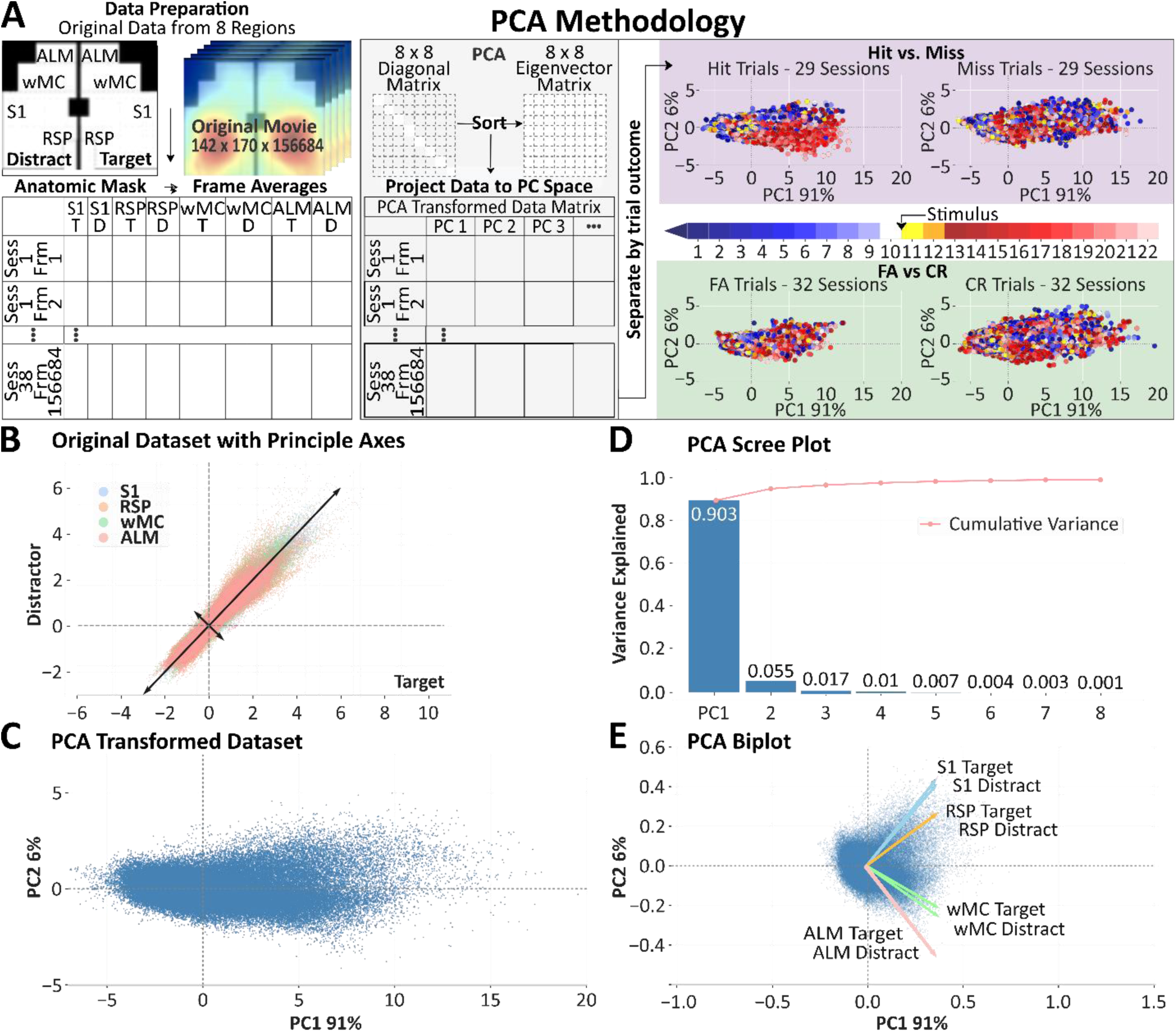
Spatial dimensionality reduction for single trial analyses. (A) Methodology for using principal component analysis (PCA) to reduce spatial dimensionality. Left, full images were parsed into 8 regional masks. Average dF/F within each mask for all trials and all sessions were appended into a single matrix, upon which PCA was performed. Right, frames with different trial outcomes were back-projected to the first principal component (PC1) and plotted against their projection onto the second principal component (PC2). Transformed samples are colored based on their frame index: prestimulus (blue to white), post-stimulus and pre-response (yellow and orange), response (red to pink). (B) Original dataset, each data point represents a sample frame ROI-specific average, plotted against its change in fluorescence (dF/F) between target (x-axis) and distractor (y-axis) hemispheres. Black arrows represent the first two principal vectors. (C) Transformed dataset, each data point represents a sample frame plotted against its projection onto PC1 and PC2. (D) PCA scree plot. PCs are plotted according to their rank in variance, with accumulated variance plotted in red. The first two PCs were chosen for further analysis as they explain >95% variance of the untransformed dataset (PC1, 91%, PC2, 6%). (E) PCA biplot. Samples plotted against their normalized projection onto PC1 and PC2, with vectors representing individual ROIs according to their loadings.

We determined the distributions of prestimulus activity from single frames within this PCA space (Figure 7). In Figure 7A, we plot the data from two example sessions, in which each data point is a single prestimulus frame preceding a hit (purple) or miss (yellow) trials. We noticed that the data from hit trials were more tightly clustered than the data from miss trials. To quantify this observation, first we fit the data from each trial type with a covariance ellipse. The shaded ellipses in Figure 7A represent a confidence area of 1 standard deviation, σ, which we used as a measure of framewise variability. Figure 7B plots the confidence area for prestimulus activity on hit and miss trials for all sessions (n=29 sessions). The prestimulus activity variance is significantly lower for hit compared to miss trials (effect size, Cohen’s d=1.92; paired t-test, t(28)= 9.43, p=1.74e-10).

**Figure 7:**
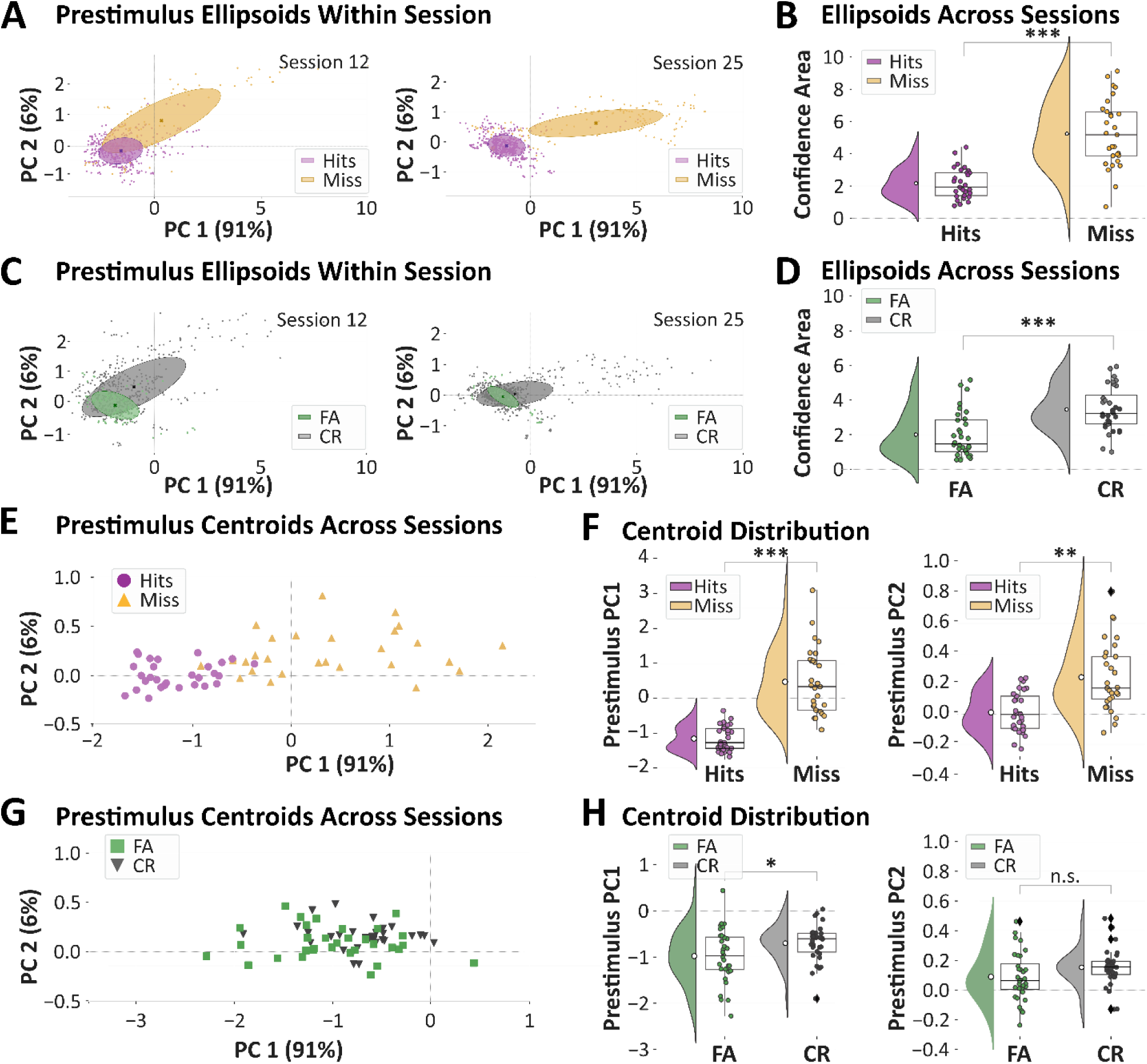
Single trial analyses of prestimulus subspace variance and position according to trial outcomes. All data presented are from the last 500 ms of the prestimulus window (frames 6 to 10 of Figure 6A). (A) Prestimulus activity in PC space for hit (purple) and miss (yellow) trials of two example sessions. Each data point represents a single prestimulus frame. Overlaid are covariance ellipses for both trial outcome types (major radius, 1σ along PC1; minor radius, 1σ along PC2). Note the reduced area and distinct position of the covariance ellipses for hit compared to miss trials. (B) Comparison of the ellipse area, as a measure of variability, across all sessions. (C and D) Same as [A] and [B], except for FA (green) and CR (gray) trials. Response trials (hit and FA) are preceded by less variable prestimulus activity compared to no response trials (miss and CR). (E) Centroid positions of the covariance ellipses in PC space for all sessions, for hit and miss trials (same color designation as above). Each data point represents the hit or miss centroid from one session. (F) Quantification of centroid positions on axes PC1 (left) and PC2 (right). (G and H) Same as [E] and [F], except for FA and CR trials. Prestimulus activity occupies distinct subspaces for response and no response trials, along both PC1 and PC2 for target trials and along PC 1 for distractor trials. *p<0. 01; **p<0.001; ***p<0.0001; n.s., non-significant.

We conducted the same analyses for distractor trials and obtained similar results. The two example sessions in Figure 7C show more tightly clustered prestimulus activity for response (FA) compared to no response (CR) trials. Across all sessions (n=32), the confidence areas are significantly lower for FA compared to CR trials (effect size, Cohen’s d=1.11; paired t-test, t(31)= 7.40, p=1.22e-8, Figure 7D). Thus, for both target and distractor trials, lower framewise prestimulus variability predicts response outcomes.

In addition to differences in variability, we also noticed that the prestimulus activity resides in different subspaces preceding response and no response trials. As evident in Figure 7A, *within* each session the centroids of the hit and miss confidence areas are offset, whereas *between* these two sessions the hit centroids occur at similar positions. In Figure 7E, we plot the centroid position for all sessions (n=29 sessions). Indeed, we find that across all sessions the centroid positions preceding hit trials are separated from the centroid positions preceding miss trials. This separation is significant, for both PC1 and PC2 axes (Figure 7F, PC1: d=2.19, paired t-test, t(28)=8.55, p=1.34e-9; PC2: d=1.24, t(28)=4.01, p=2.07e-4). In contrast, for distractor trials, the centroids of prestimulus activity show greater overlap for response (FA) and no response (CR) trials (Figure 7G). However, we do still find significantly different centroid positions on distractor trials along PC1 (Figure 7H, PC1: d=0.57, paired t-test, t(31)=2.99, p=0.0027; PC2: d=0.43, t(31)=1.30, p=0.10). These data indicate that the neuronal activity across dorsal neocortex preceding response trials is less variable than no response trials and occupies a separate subspace. Similar to prestimulus neural activity (Figure 3) and movement (Figure 5), the differences in variability and subspace position are larger for target compared to distractor trials. Taken together, these data specify an optimal neuronal and behavioral state for stimulus detection.

### Distribution of prestimulus choice probability among single units

The above analyses of widefield imaging data assessed population neuronal activity. In this final series of analyses, we sought to determine the distribution of task-relevant prestimulus activities among single units (Figure 8). During the same selective whisker detection task, we recorded 936 single units, from target-aligned S1 (377 units), target-aligned wMC (338 units) and target-aligned ALM (221 units). First, we quantified the prestimulus choice probability of all units on target trials. Choice probabilities (CP) of single units in each region were marginally below chance (Figure 8A, CP μ_S1_=49.10 ± 0.16, one-sample t-test, t(376)=-5.72, p=2.21e-8, CP μ_wMC_=49.44 ±0.2, one-sample t-test, t(337)=-2.82, p=0.005, CP μ_ALM_=49.64 ±0.19, one-sample t-test, t(220)=-1.92, p=0.06). These distributions were not significantly different across the three regions (two-way ANOVA: F(2,933)= 2.18, p=0.11 and post hoc Tukey: S1 vs. wMC, p=0.33; wMC vs. ALM, p=0.76; S1 vs. ALM, p=0.12). Prestimulus choice probability below chance indicates that lower activity predicts hit compared to miss outcomes, and therefore is consistent with the widefield imaging data. However, the distributions of these data indicate that only a small portion of single units show strong prestimulus choice probability.

**Figure 8:**
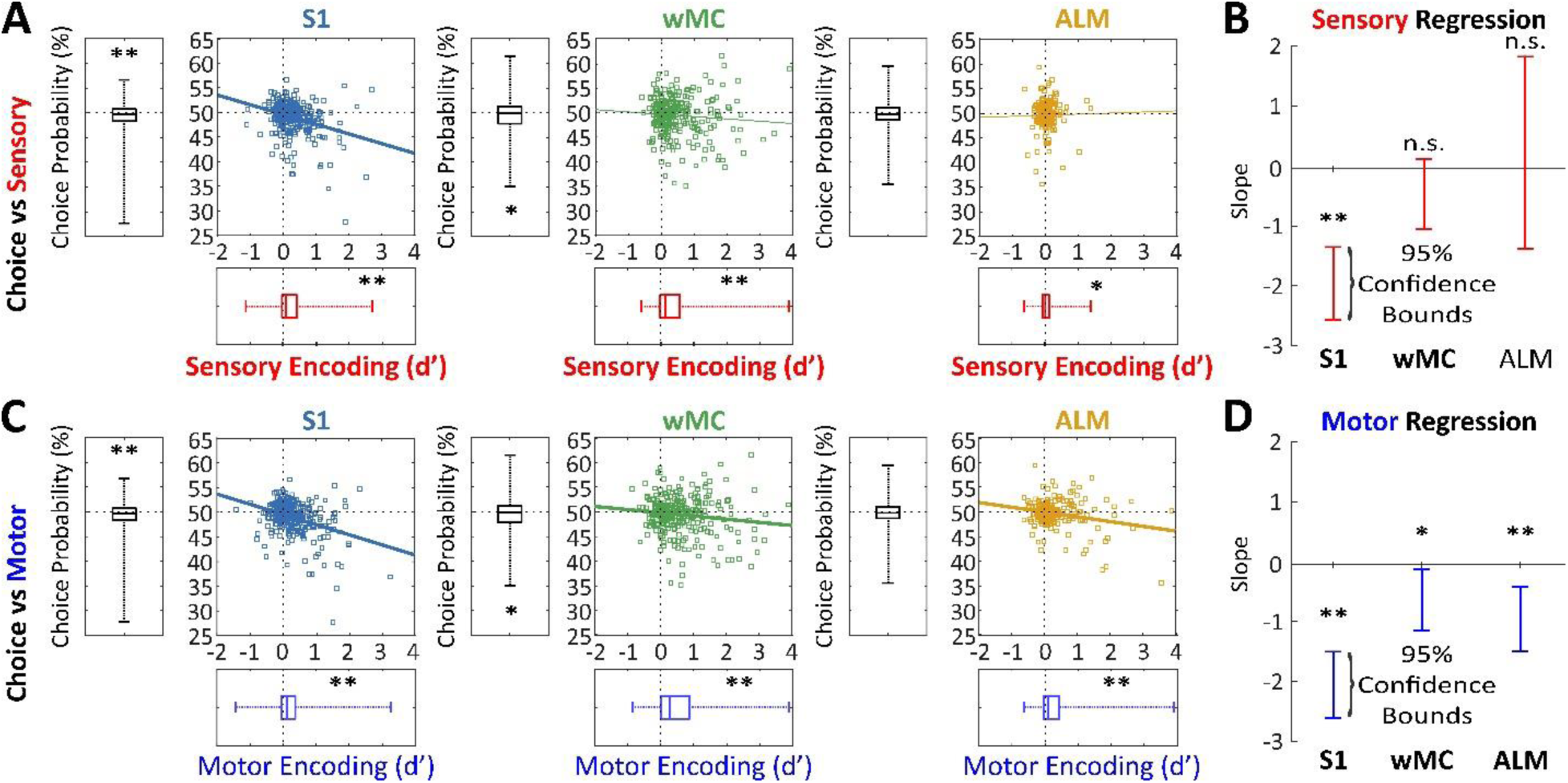
Distribution of prestimulus choice probability, post-stimulus sensory, and pre-response motor encoding across single units in S1, wMC and ALM. (A) Plots of sensory encoding (d’) versus choice probability (%) for single units in target-aligned S1 (left), wMC (center), and ALM (right). Asterisks above box plots reflect comparisons of individual measures to chance (d’=0 and choice probability=50%). Scatter plots include linear fits of the single unit data. Single units in each of these three cortical regions show below chance prestimulus choice probability (tending yet not significant for ALM (p=0.06), significant for S1 and wMC) and positive post-stimulus sensory encoding. (B) 95% confidence bounds of the linear regression slope values. (C and D) Same as [A] and [B], but for pre-response motor encoding. The significant negative slope values indicate an overlap between the single units with lower than chance prestimulus choice probability and positive post-stimulus sensory encoding (for S1) and pre-response motor encoding (for S1, wMC, and ALM). *p<0.05; **p<0.005; n.s., non-significant.

Given this variability of single units, we next asked whether the units with strong prestimulus choice probability overlap with the units with strong post-stimulus sensory and pre-response motor encoding. To test this, we plotted prestimulus choice probability against post-stimulus sensory (Figure 8A) and pre-response motor (Figure 8B) encoding. The negative regression slopes show correlations between choice probability and sensory encoding for S1, and between choice probability and motor encoding for S1, wMC, and ALM (Figure 8C and 8D, one-sample t-test, sensory encoding slope: m_S1_=-1.96 ± 0.31, t(375)=-6.34, p=6.56e-10; one-sample t-test, motor encoding slope: m_S1_=-2.05 ± 0.28, t(375)=-7.35, p=1.23e-12, m_wMC_=-0.64 ± 0.26, t(336)=-2.49, p=0.013, m_ALM_=-0.96 ± 0.28, t(219)=-3.46, p=6.41e-4). Thus, units in these regions have combined neuronal representations such that those representing prestimulus behavior context overlap with those with post-stimulus (sensory) and pre-response (motor) task-relevant encoding. This overlap may be influenced by a common factor such as firing rate (Supplemental Figure 2). Nevertheless, these analyses demonstrate that the subset of neurons that show the largest prestimulus suppression on hit trials are the same neurons that strongly encode task features.

## Discussion

The primary focus of this study is to determine whether and how neuronal activity before stimulus onset predicts trial outcomes during goal-directed behavior. We assessed this for both target and distractor stimulus detection. We find that lower prestimulus activity predicts detection of both target and distractor stimuli (Figures 2 and 3) and faster reaction times on Hit trials (Figure 3). This low activity state is distributed globally throughout dorsal cortex (Figure 3), maps onto a distinct, less variable subspace than activity preceding no response trials (Figure 7) and is represented most robustly in the subset of neurons also encoding post-stimulus sensory and pre-response motor task features (Figure 8).

The impacts of spontaneous activity on stimulus responses have been explored extensively in both physiological and computational studies. Increased spontaneous activity has been proposed to increase response gain by two primary mechanisms: depolarization to reduce membrane potential distance to spike threshold and increased variance to amplify the impacts of weak inputs (Cardin et al., 2008; Haider et al., 2007; Haider & McCormick, 2009; Hô & Destexhe, 2000; Rudolph & Destexhe, 2003; Shu et al., 2003). Therefore, we were surprised to find that reduced prestimulus activity correlated with both enhanced stimulus detection (Figures 2 and 3) and increased sensory responses (Figure 4). And yet, our data are consistent with studies in primary somatosensory and auditory cortices, demonstrating increased sensory responses with reduced prestimulus activity (Hasenstaub et al., 2007; McGinley, David, et al., 2015; Sachdev et al., 2004). Future studies are required to determine the cellular and network mechanisms underlying increased responsiveness with low activity, with possibilities including reduced membrane conductance (Chance et al., 2002), reduced inhibition, and reduced synaptic depression.

Our study was conducted in the context of a somatosensory (whisker) detection task. It is not currently known, however, whether these findings will generalize to other sensory modalities and other types of tasks. Reduced network activity and reduced synaptic variance have been shown to predict a network with a discrete, all-or-none input-output function (Hô & Destexhe, 2000). This configuration may improve distinguishing the presence versus absence of a stimulus as needed for stimulus detection. Such a network state, though, would be predicted to poorly encode the precise features of a stimulus. Therefore, we speculate that tasks requiring discrimination of fine stimulus details may be optimal in a high activity network state with a continuous input-output function. However, this remains to be tested.

Most studies of the impacts of spontaneous activity on sensory responses focus on primary sensory areas. However, stimulus detection tasks require the contributions of multiple cortices (de Lafuente and Romo 2006). Indeed, we have recently shown that the task in this study activates multiple sensory and motor cortices, including S1, wMC, and ALM (Aruljothi et al., 2020; Zareian et al., 2021). In this study we demonstrate that the prestimulus activity predictive of trial outcome is global, involving all regions of dorsal neocortex. This global cortical state may reflect the coordination amongst multiple cortices, to improve not just stimulus encoding in primary sensory cortex, but the propagation of task-relevant signals throughout neocortex. Interestingly, we found prestimulus activity suppression to be largest in the same neurons that also strongly encode post-stimulus sensory and pre-response motor features, in S1, wMC, and ALM. This organization may ensure coordination not just between cortical regions, but among the specific neuronal ensembles involved in this stimulus detection task. Low activity in these specific neuronal ensembles may increase excitability and transmission by increasing membrane resistance and reducing synaptic depression.

Global changes in cortical state, as observed here, are traditionally associated with changes in arousal, driven by widespread ascending neuromodulatory systems (Zagha & McCormick, 2014). More recently, studies have shown that movement is associated with global increases in neocortical activity (Musall et al., 2020; Salkoff et al., 2020; Stringer et al., 2019). As with low activity preceding response trials, we also find that whisker movements are reduced preceding hit trials (Figure 5), consistent with previous reports (Kyriakatos et al., 2016; Ollerenshaw et al., 2012). We suspect that whisker movements impair detection for multiple reasons: 1) reafference signals from self-generated movements (Fee et al., 1997) may obscure stimulus-evoked afferent signals, 2) self-generated movements may evoke top-down sensory gating and thereby suppress stimulus evoked signals (Chakrabarti & Schwarz, 2018), and 3) centrally-mediated cortical activation associated with whisker movements (Poulet et al., 2012) may reduce network excitability. And yet, our findings support a view of cortical state as higher dimensional than stationary versus moving (Zagha and McCormick 2014; McGinley, Vinck et al., 2015). Among Hit trials, we find a positive correlation between prestimulus activity and reaction time (Figure 3). This suggests that even within overt changes in arousal, the precise levels of cortical activity impact performance in our task, with the lowest prestimulus activity correlating with optimal performance. Dissecting the physiological mechanisms underlying the low amplitude cortical state permissive for whisker stimulus detection is a focus of ongoing investigations.

## Acknowledgements

This work was supported by the Whitehall Foundation (Research Grant 2017-05-71 to E.Z.) and the National Institutes of Health (R01NS107599 to E.Z.). We thank Drs. Hongdian Yang and David Salkoff for many helpful discussions throughout the project. The authors declare no competing financial interests.

## Figures and Legends

**Supplemental Figure 1.**
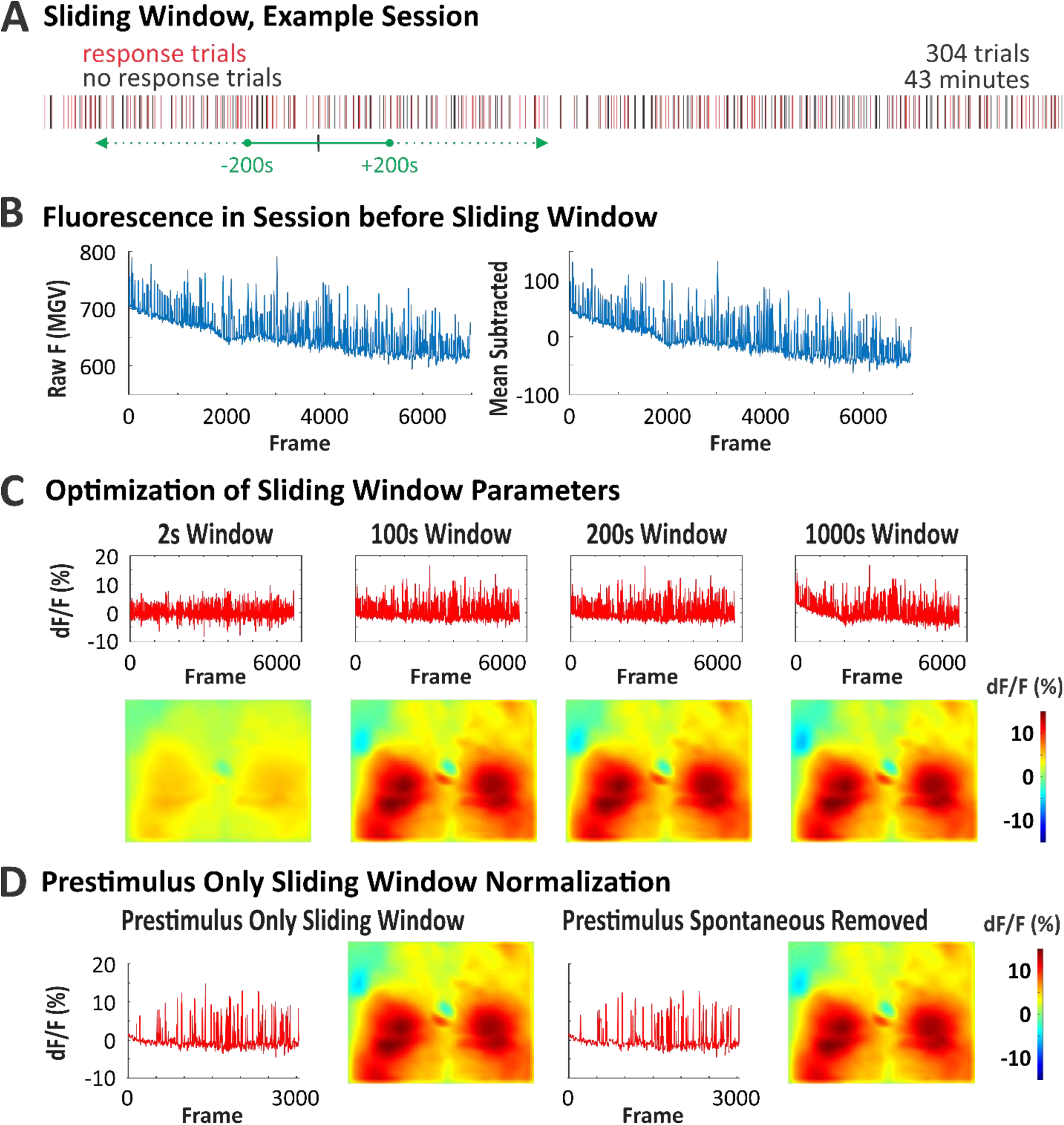
Related to Methods, Figures 1-3: Sliding window normalization method and robustness of window size. (A) Depiction of response trials (red), no response trials (black) and sliding window (green) used for an example session. This session consisted of 304 trials over 43.4 minutes. Each sliding window segment included an average of 46 trials. (B) Raw fluorescence per frame and mean subtracted raw fluorescence per frame acquired across example session. Rundown throughout the recording session is not corrected by mean subtraction. (C) Different sliding windows considered for optimization of method used in this study. Top row: dF/F using a sliding window every 2s; 2s half-width (far left), 100s half-width (center left), 200s half-width (center right), 1000s half-width (far right). Bottom row: Miss to Hits difference using the sliding window indicated in top row. This normalization method is robust to a range of sliding window sizes, between 50s to 200s. If the window is too small (left) single trial differences are normalized out. If the window is too large (right) fluorescence rundown is not corrected. (D) Sliding window method (200s) applied to prestimulus frames only and applied to prestimulus frames with spontaneous trials removed. Left to right: dF/F per frame across prestimulus frames in example session (far left), Miss to Hits difference using only prestimulus frames (left center), dF/F per frame across prestimulus frames, spontaneous trials removed, in example session (right center), Miss to Hits difference using prestimulus frames, spontaneous trials removed (far right). Excluding post-stimulus frames and spontaneous trials does not impact our sliding window prestimulus analyses.

**Supplemental Figure 2.**
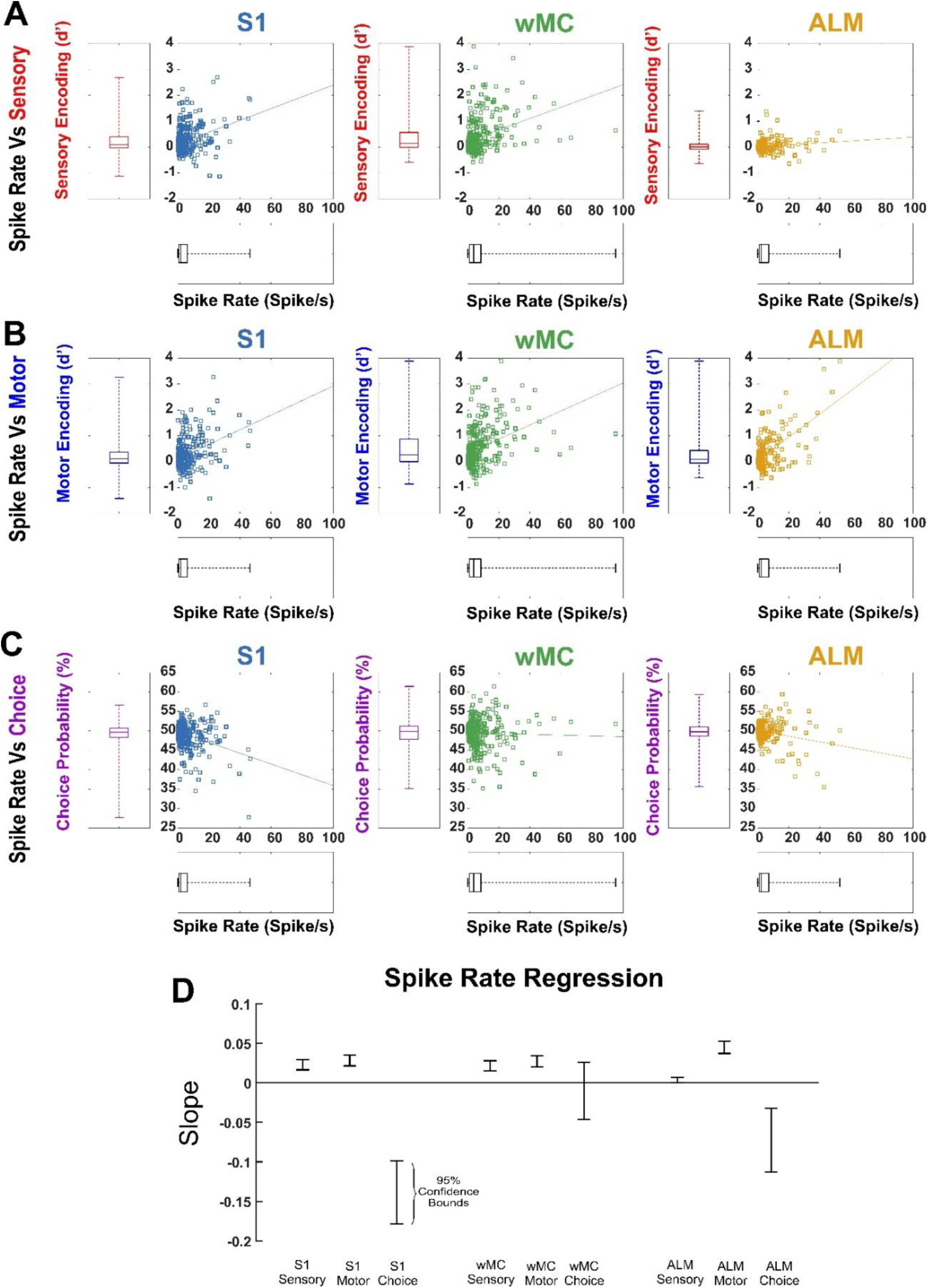
Related to Figure 8: Relationship between spike rate and prestimulus choice probability and post-stimulus sensory and pre-response motor encoding across single units in S1, wMC and ALM. (A) Plots of sensory encoding (d’) versus spike rate (Hz) for single units in target-aligned S1 (left), wMC (center), and ALM (right). Scatter plots include linear fits of the single unit data. Single units in each of these three cortical regions show positive relationship between spike rate and post-stimulus sensory encoding. (B and C) Same as [A], but for pre-response motor encoding (B) and prestimulus choice probability (C). (D) 95% confidence bounds of the linear regression slope values for all scatter plots. These data identify spike rate as a common factor that correlates with both post-stimulus sensory and motor encoding (positive correlation) and prestimulus choice probability (negative correlation).

## References

Andreou, C., & Borgwardt, S. (2020). Structural and functional imaging markers for susceptibility to psychosis. Mol Psychiatry, 25(11), 2773–2785. https://doi.org/10.1038/s41380-020-0679-7

Arieli, A., Shoham, D., Hildesheim, R., & Grinvald, A. (1995). Coherent spatiotemporal patterns of ongoing activity revealed by real-time optical imaging coupled with single-unit recording in the cat visual cortex. Journal of Neurophysiology, 73(5), 2072–2093. https://doi.org/10.1152/jn.1995.73.5.2072

Arieli, A., Sterkin, A., Grinvald, A., & Aertsen, A. (1996). Dynamics of Ongoing Activity: Explanation of the Large Variability in Evoked Cortical Responses. Science, 273(5283), 1868–1871. https://doi.org/10.1126/science.273.5283.1868

Aruljothi, K., Marrero, K., Zhang, Z., Zareian, B., & Zagha, E. (2020). Functional Localization of an Attenuating Filter within Cortex for a Selective Detection Task in Mice. The Journal of Neuroscience, 40(28), 5443–5454. https://doi.org/10.1523/jneurosci.2993-19.2020

Boly, M., Balteau, E., Schnakers, C., Degueldre, C., Moonen, G., Luxen, A., Phillips, C., Peigneux, P., Maquet, P., & Laureys, S. (2007). Baseline brain activity fluctuations predict somatosensory perception in humans. Proceedings of the National Academy of Sciences, 104(29), 12187–12192. https://doi.org/10.1073/pnas.0611404104

Britten, K., Shadlen, M., Newsome, W., & Movshon, J. (1992). The analysis of visual motion: a comparison of neuronal and psychophysical performance. The Journal of Neuroscience, 12(12), 4745–4765. https://doi.org/10.1523/jneurosci.12-12-04745.1992

Cardin, J. A., Palmer, L. A., & Contreras, D. (2008). Cellular Mechanisms Underlying Stimulus-Dependent Gain Modulation in Primary Visual Cortex Neurons In Vivo. Neuron, 59(1), 150–160. https://doi.org/10.1016/j.neuron.2008.05.002

Chakrabarti, S., & Schwarz, C. (2018). Cortical modulation of sensory flow during active touch in the rat whisker system. Nature Communications, 9(1). https://doi.org/10.1038/s41467-018-06200-6

Chance, F. S., Abbott, L. F., & Reyes, A. D. (2002). Gain Modulation from Background Synaptic Input. Neuron, 35(4), 773–782. https://doi.org/10.1016/s0896-6273(02)00820-6

Crochet, S., & Petersen, C. C. (2006). Correlating whisker behavior with membrane potential in barrel cortex of awake mice. Nat Neurosci, 9(5), 608–610. https://doi.org/10.1038/nn1690

de Lange, F. P., Rahnev, D. A., Donner, T. H., & Lau, H. (2013). Prestimulus oscillatory activity over motor cortex reflects perceptual expectations. J Neurosci, 33(4), 1400–1410. https://doi.org/10.1523/JNEUROSCI.1094-12.2013

Fee, M. S., Mitra, P. P., & Kleinfeld, D. (1997). Central Versus Peripheral Determinants of Patterned Spike Activity in Rat Vibrissa Cortex During Whisking. Journal of Neurophysiology, 78(2), 1144–1149. https://doi.org/10.1152/jn.1997.78.2.1144

Ferezou, I., Haiss, F., Gentet, L. J., Aronoff, R., Weber, B., & Petersen, C. C. (2007). Spatiotemporal dynamics of cortical sensorimotor integration in behaving mice. Neuron, 56(5), 907–923. https://doi.org/10.1016/j.neuron.2007.10.007

Fiebelkorn, I. C., & Kastner, S. (2021). Spike Timing in the Attention Network Predicts Behavioral Outcome Prior to Target Selection. Neuron, 109(1), 177–188 e174. https://doi.org/10.1016/j.neuron.2020.09.039

Fries, P., Reynolds, J. H., Rorie, A. E., & Desimone, R. (2001). Modulation of Oscillatory Neuronal Synchronization by Selective Visual Attention. Science, 291(5508), 1560–1563. https://doi.org/10.1126/science.1055465

Ghose, G. M., & Maunsell, J. H. R. (2002). Attentional modulation in visual cortex depends on task timing. Nature, 419(6907), 616–620. https://doi.org/10.1038/nature01057

Gold, J. I., & Shadlen, M. N. (2007). The neural basis of decision making. Annu Rev Neurosci, 30, 535–574. https://doi.org/10.1146/annurev.neuro.29.051605.113038

Haider, B., Duque, A., Hasenstaub, A. R., Yu, Y., & McCormick, D. A. (2007). Enhancement of Visual Responsiveness by Spontaneous Local Network Activity In Vivo. Journal of Neurophysiology, 97(6), 4186–4202. https://doi.org/10.1152/jn.01114.2006

Haider, B., & McCormick, D. A. (2009). Rapid neocortical dynamics: cellular and network mechanisms. Neuron, 62(2), 171–189. https://doi.org/10.1016/j.neuron.2009.04.008

Hanes, D. P., & Schall, J. D. (1996). Neural Control of Voluntary Movement Initiation. Science, 274(5286), 427–430. https://doi.org/10.1126/science.274.5286.427

Hasenstaub, A., Sachdev, R. N. S., & McCormick, D. A. (2007). State Changes Rapidly Modulate Cortical Neuronal Responsiveness. Journal of Neuroscience, 27(36), 9607–9622. https://doi.org/10.1523/jneurosci.2184-07.2007

Hô, N., & Destexhe, A. (2000). Synaptic Background Activity Enhances the Responsiveness of Neocortical Pyramidal Neurons. Journal of Neurophysiology, 84(3), 1488–1496. https://doi.org/10.1152/jn.2000.84.3.1488

Kim, R., & Sejnowski, T. J. (2021). Strong inhibitory signaling underlies stable temporal dynamics and working memory in spiking neural networks. Nat Neurosci, 24(1), 129–139. https://doi.org/10.1038/s41593-020-00753-w

Kyriakatos, A., Sadashivaiah, V., Zhang, Y., Motta, A., Auffret, M., & Petersen, C. C. H. (2016). Voltage-sensitive dye imaging of mouse neocortex during a whisker detection task. Neurophotonics, 4(3), 031204. https://doi.org/10.1117/1.NPh.4.3.031204

Luck, S. J., Chelazzi, L., Hillyard, S. A., & Desimone, R. (1997). Neural Mechanisms of Spatial Selective Attention in Areas V1, V2, and V4 of Macaque Visual Cortex. Journal of Neurophysiology, 77(1), 24–42. https://doi.org/10.1152/jn.1997.77.1.24

Mazaheri, A., DiQuattro, N. E., Bengson, J., & Geng, J. J. (2011). Pre-stimulus activity predicts the winner of top-down vs. bottom-up attentional selection. PLoS One, 6(2), e16243. https://doi.org/10.1371/journal.pone.0016243

McCormick, D. A., McGinley, M. J., & Salkoff, D. B. (2015). Brain state dependent activity in the cortex and thalamus. Curr Opin Neurobiol, 31, 133–140. https://doi.org/10.1016/j.conb.2014.10.003

McGinley, M. J., David, S. V., & McCormick, D. A. (2015). Cortical Membrane Potential Signature of Optimal States for Sensory Signal Detection. Neuron, 87(1), 179–192. https://doi.org/10.1016/j.neuron.2015.05.038

McGinley, M. J., Vinck, M., Reimer, J., Batista-Brito, R., Zagha, E., Cadwell, C. R., Tolias, A. S., Cardin, J. A., & McCormick, D. A. (2015). Waking State: Rapid Variations Modulate Neural and Behavioral Responses. Neuron, 87(6), 1143–1161. https://doi.org/10.1016/j.neuron.2015.09.012

Moore, T., & Armstrong, K. M. (2003). Selective gating of visual signals by microstimulation of frontal cortex. Nature, 421(6921), 370–373. https://doi.org/10.1038/nature01341

Murphy, M. C., Chan, K. C., Kim, S. G., & Vazquez, A. L. (2018). Macroscale variation in resting-state neuronal activity and connectivity assessed by simultaneous calcium imaging, hemodynamic imaging and electrophysiology. Neuroimage, 169, 352–362. https://doi.org/10.1016/j.neuroimage.2017.12.070

Musall, S., Kaufman, M. T., Juavinett, A. L., Gluf, S., & Churchland, A. K. (2020). Single-trial neural dynamics are dominated by richly varied movements. Nat Neurosci, 22(10), 1677–1686. https://doi.org/10.1038/s41593-019-0502-4

Niell, C. M., & Stryker, M. P. (2010). Modulation of visual responses by behavioral state in mouse visual cortex. Neuron, 65(4), 472–479. https://doi.org/10.1016/j.neuron.2010.01.033

Ollerenshaw, D. R., Bari, B. A., Millard, D. C., Orr, L. E., Wang, Q., & Stanley, G. B. (2012). Detection of tactile inputs in the rat vibrissa pathway. Journal of Neurophysiology, 108(2), 479–490. https://doi.org/10.1152/jn.00004.2012

Petersen, C. C. H., Grinvald, A., & Sakmann, B. (2003). Spatiotemporal Dynamics of Sensory Responses in Layer 2/3 of Rat Barrel Cortex MeasuredIn Vivoby Voltage-Sensitive Dye Imaging Combined with Whole-Cell Voltage Recordings and Neuron Reconstructions. The Journal of Neuroscience, 23(4), 1298–1309. https://doi.org/10.1523/jneurosci.23-04-01298.2003

Poulet, J. F., Fernandez, L. M., Crochet, S., & Petersen, C. C. (2012). Thalamic control of cortical states. Nat Neurosci, 15(3), 370–372. https://doi.org/10.1038/nn.3035

Poulet, J. F., & Petersen, C. C. (2008). Internal brain state regulates membrane potential synchrony in barrel cortex of behaving mice. Nature, 454(7206), 881–885. https://doi.org/10.1038/nature07150

Roitman, J. D., & Shadlen, M. N. (2002). Response of Neurons in the Lateral Intraparietal Area during a Combined Visual Discrimination Reaction Time Task. The Journal of Neuroscience, 22(21), 9475–9489. https://doi.org/10.1523/jneurosci.22-21-09475.2002

Rudolph, M., & Destexhe, A. (2003). A Fast-Conducting, Stochastic Integrative Mode for Neocortical NeuronsInVivo. The Journal of Neuroscience, 23(6), 2466–2476. https://doi.org/10.1523/jneurosci.23-06-02466.2003

Sachdev, R. N. S., Ebner, F. F., & Wilson, C. J. (2004). Effect of Subthreshold Up and Down States on the Whisker-Evoked Response in Somatosensory Cortex. Journal of Neurophysiology, 92(6), 3511–3521. https://doi.org/10.1152/jn.00347.2004

Sachidhanandam, S., Sreenivasan, V., Kyriakatos, A., Kremer, Y., & Petersen, C. C. H. (2013). Membrane potential correlates of sensory perception in mouse barrel cortex. Nature Neuroscience, 16(11), 1671–1677. https://doi.org/10.1038/nn.3532

Salkoff, D. B., Zagha, E., McCarthy, E., & McCormick, D. A. (2020). Movement and Performance Explain Widespread Cortical Activity in a Visual Detection Task. Cereb Cortex, 30(1), 421–437. https://doi.org/10.1093/cercor/bhz206

Shimaoka, D., Harris, K. D., & Carandini, M. (2018). Effects of Arousal on Mouse Sensory Cortex Depend on Modality. Cell Reports, 22(12), 3160–3167. https://doi.org/10.1016/j.celrep.2018.02.092

Shu, Y., Hasenstaub, A., Badoual, M., Bal, T., & McCormick, D. A. (2003). Barrages of Synaptic Activity Control the Gain and Sensitivity of Cortical Neurons. The Journal of Neuroscience, 23(32), 10388–10401. https://doi.org/10.1523/jneurosci.23-32-10388.2003

Stringer, C., Pachitariu, M., Steinmetz, N., Reddy, C. B., Carandini, M., & Harris, K. D. (2019). Spontaneous behaviors drive multidimensional, brainwide activity. Science, 364(6437), 255. https://doi.org/10.1126/science.aav7893

van Kempen, J., Gieselmann, M. A., Boyd, M., Steinmetz, N. A., Moore, T., Engel, T. A., & Thiele, A. (2020). Top-down coordination of local cortical state during selective attention. Neuron. https://doi.org/10.1016/j.neuron.2020.12.013

Yang, H., Kwon, S. E., Severson, K. S., & O’Connor, D. H. (2016). Origins of choice-related activity in mouse somatosensory cortex. Nat Neurosci, 19(1), 127–134. https://doi.org/10.1038/nn.4183

Zagha, E., & McCormick, D. A. (2014). Neural control of brain state. Curr Opin Neurobiol, 29, 178–186. https://doi.org/10.1016/j.conb.2014.09.010

Zareian, B., Maboudi, K., Daliri, M. R., Abrishami Moghaddam, H., Treue, S., & Esghaei, M. (2020). Attention strengthens across-trial pre-stimulus phase coherence in visual cortex, enhancing stimulus processing. Sci Rep, 10(1), 4837. https://doi.org/10.1038/s41598-020-61359-7

Zareian, B., Zhang, Z., & Zagha, E. (2021). Cortical Localization of the Sensory-Motor Transformation in a Whisker Detection Task in Mice. eNeuro, ENEURO.0004-0021. https://doi.org/10.1523/eneuro.0004-21.2021

